# Relative biological effect of alpha particle radiation on low dose phenomena: lethal mutation, hyper-radiosensitivity and increased radioresistance

**DOI:** 10.1101/435487

**Authors:** Chandula Fernando, Xiaopei Shi, Soo Hyun Byun, Colin B. Seymour, Carmel E. Mothersill

## Abstract

At high doses, the current recommended radiation weighting factors advise a significantly higher effectiveness of alpha particles relative to gamma radiation. However, at lower doses, the ratio of effectiveness between radiations of varying linear energy transfer values is complicated due to the relative importance of low dose phenomena such as genomic instability, bystander effects, low dose hyper-radiosensitivity and increased radioresistance (HRS/IRR). Radium is the most common source of alpha radiation exposure to humans, but the dosimetry is complicated by the decay chain which involves gamma exposure due to radon daughters. This study aimed to isolate the relative biological effect of alpha particles after low doses of radium to cells and their progeny. This was done by subtracting the survival values of a human keratinocyte cell line (HaCaT) and an embryonic Chinook salmon cell line (CHSE-214) exposed to gamma irradiation, from survival of the same cell lines exposed to mixed alpha and gamma radiation through chronic exposure to Ra-226 and its decay products. The human cell line showed increased radioresistance when exposed to low doses of alpha particles. In contrast the fish cell line, which demonstrated radioresistance to low dose gamma energy, demonstrated increased lethality when exposed to low doses of alpha particles. The results confirm the need to consider the dose-response relationship when developing radiation weighting factors for low dose exposures, as well as the need to be aware of possible cell line and species differences.

## Introduction

Linear energy transfer (LET), describes the amount of energy deposited to the interacting material, per unit of distance. Photons such as gamma rays are able to traverse great distances unchanged before being absorbed, however monoenergetic ions such as alpha particles cause frequent direct ionizations within a smaller range. Due in part to the clustered nature of damage caused, the relative biological effectiveness (RBE) of an alpha particle is often described to be significantly higher than that of a gamma ray. This is a result of the concentration of damage given the same amount of absorbed energy [1].

While this may be true for high doses, it has been shown that if a single alpha particle traverses a cell, it causes zero to small risk of oncogenic transformation [2]. Further, the work of Nagasawa and Little has shown significantly higher frequencies of mutation than would be expected through linear extrapolation from data for high doses, at doses where the mean number of alpha particle traversals per nucleus was significantly less than one [3]. At low doses, both alpha and gamma rays can cause non-targeted effects (NTE) like genomic instability where damage does not cause direct mortality and cells appear completely normal but de novo effects are seen in distant progeny and lethality occurs generations later (often referred to as lethal mutations or delayed reproductive death) [4]. Cell survival at sub-lethal doses of gamma irradiation has also been observed to differ from what is expected by the traditional linear-quadratic model, instead displaying a region of low-dose hyper-radiosensitivity (HRS) followed by increased radioresistance (IRR) [5]. Currently accepted recommendations for the radiation weighting factor (w_r) of alpha particles, which apply the concept of RBE to derive equivalent dose, are dose independent [6]. However, research has shown instances for RBE to be dose dependent when high dose biological effects are substantially different to low dose effects [7].

Of particular interest to this study is whether NTE amplify low dose effects such that they are higher than what would be expected from established linear no-threshold model (LNT) related RBE values following exposure to low doses of an environmental alpha emitter: radium-226. Despite being an alpha emitter by itself, it is known that the uranium decay chain of which radium is part of involves many gamma emissions, thereby making it difficult to measure pure alpha effects. To approach this problem, observations from gamma irradiation (through acute exposure to Cs-137) will be subtracted from mixed alpha and gamma irradiation (through chronic exposure to Ra-226 and its progeny).

This study will measure the acute survival and the lethal mutation phenotype assayed as reduced cloning efficiency in culture of a human keratinocyte cell line (HaCaT). In addition, due to the increasing relevance of protecting non-human biota from radium in hydrogeologic contaminations from mining, etc., this study will also investigate relative alpha exposure effects in the embryonic Chinook salmon cell line (CHSE-214).

## Materials and Methods

### Cell culture

The HaCaT cell line used in the study is an immortalized human keratinocyte cell line originally derived and characterized by Boukamp et al [8]. The cell line used in this study was obtained as a gift from Dr. Orla Howe (Dublin, Ireland). The cell line was routinely maintained with RPMI-1640 medium supplemented with 10% fetal bovine serum (Invitrogen, Burlington, Canada), 5 ml of 200 mM L-Glutamine (Gibco, Burlington, Canada), 0.5 g/ml hydrocortisone (Sigma-Aldrich, Oakville, Canada), 25 mM Hepes buffer (Gibco), penicillin and streptomycin (Gibco). These cells were grown at 37°C in an incubator with 5% CO_2_.

The CHSE-214 is an embryonic cell line derived from Chinook salmon obtained as a gift from Dr. Neils Bols (Waterloo, Canada). CHSE-214 cells were cultured in Leibovitz’s L-15 medium supplemented with 12% fetal bovine serum (Invitrogen), 5 ml of 200 mM L-Glutamine (Gibco), 25 mM Hepes buffer (Gibco), penicillin and streptomycin (Gibco). These cells were grown at 19°C in an incubator without CO_2_.

Reduction in cloning efficiency was observed using the clonogenic assay technique developed by Puck and Marcus [9]. Cell stocks were maintained in T75 flasks with 30ml medium. Upon reaching 80-90% confluence, flasks were subcultured. Here cells were gently rinsed with calcium and magnesium-free DPBS in a biosafety level 2 laminar flow cabinet. HaCaT cells were detached using a 0.25% (v/v) trypsin-1 mM EDTA solution (Gibco) at 37°C for 8 minutes, while CHSE-214 cells were detached using a 0.125% (v/v) trypsin-1 mM EDTA solution (Gibco) at 19°C for 8 minutes. Trypsin was neutralized using fresh culture media, and the cell solution was centrifuged at 125g for 4 minutes. The pellet was resuspended, and cells were counted using an automated cell counter (Bio-Rad TC20). The cells were then seeded into fresh flasks with fresh culture media at the required cell density such that at least 100 viable colonies could be expected to form in control flasks.

Reporter T25 flasks were maintained in the incubator for 9 days. Following this incubation period, colonies in sham irradiated (control) flasks were visible to the naked eye. Flasks were stained using a 1:4 (v/v) dilution of Fuchsin-Carbol (Ricca Chemical Co., Arlington, TX) in water, and macroscopically visible colonies (confirmed to have more than 50 cells when observed under a microscope) were scored as survivors.

### Chronic irradiation using Ra-226 in medium

Stock solutions of medium containing the radioisotope Ra-226 were prepared using neutralized radium nitrate (Eckert and Ziegler, Valencia, USA). 100 ml L-15 or RPMI medium was mixed with 1000 Bq of Ra-226 solution. The concentration of Ra-226 in this stock medium was 10,000 mBq/ml. After filtering into storage tubes, serial dilutions were made to give the required final concentrations.

500 cells were initially seeded into T25 flasks containing 5 ml of medium with Ra-226 or control medium. 4 flasks were prepared for each respective concentration: 0, 0.1, 1, 10, 100, 200 or 500 mBq/ml Ra-226. Flasks were maintained in the incubator for 9 days after which the radioactive medium was removed, and the cells were gently rinsed with calcium and magnesium-free DPBS. Ra-226 residues in the flasks were assumed to be insignificant. Flasks then received 5 ml of fresh culture medium without Ra-226 and returned to the incubator. 3 flasks from each concentration were deemed reporter flasks, incubated for 9 days and stained as described above. Cloning efficiencies observed in these reporter flasks represented the initial plating efficiencies from direct chronic irradiation. The remaining fourth flask of each concentration was left to incubate until 80-90% confluency, after which it was subcultured as described above seeding 500 cells into a fresh flask. From here on however no further irradiation was to be done and all flasks received fresh culture medium containing 0 mBq/ml Ra-226. The process was repeated as before, and cloning efficiencies observed in these reporter flasks represented survival fractions of the progeny (P2). The process was repeated once more to observe further change in the cloning efficiency in subsequent generations (P3).

### Acute irradiation using a Cs-137 source

As with the Ra-226 experiments, 4 T25 flasks were seeded with 500 cells for each respective dose: 0, 0.05, 0.1, 0.25, 0.5, 0.75 or 1 Gy. The flasks were incubated for 6 hours to allow for cells to adhere to the flask, after which they were exposed to their respective γ-ray dose using a cesium-137 source (Taylor source, McMaster University, Hamilton, Canada). Flasks were placed at 26 cm from the radiation source, irradiated at a dose rate of 0.273 Gy/min and the room temperature was around 26°C.

All flasks were placed back in the incubator immediately after irradiation. Similar to the Ra-226 experiments, 3 flasks were deemed reporter flasks and incubated for approximately 9 days before being stained as described above (initial). The remaining fourth flask of each dose was incubated until cells became 80-90% confluent, after which they were subcultured as described above with fresh culture medium. This process was also repeated twice as above (P2 and P3).

### Determining γ dose from Ra-226

All possible γ emission events during the decay of Ra-226 to daughters Pb-214 and Bi-214 were tabulated according to their energy (keV) and probability (%) [10]. Total γ energy emitted per decay was then found through the summation of each γ energy multiplied by its emission probability. A system was then set up using MonteCarlito 1.10 to describe the starting source geometry: a plane of uniformly spread particles with a y-dimension twice as big as the x-dimension (similar to T-25 flask dimensions), with each particle representing emitted γ energy from the source. The radionuclide was assumed to be evenly distributed in the medium. The number of particles was calculated through the concentration of Ra-226 in each respective medium (dividing the activity by the decay constant). The average distance for one interaction by an emitted particle was calculated through its mean free path. Central particles conducted most interactions within the flask however only a quarter of interactions of particles closer to the corners of the flask, and half of interactions from particles immediately adjacent to flask walls contributed dose within the flask. Figure 1 shows one iteration of the Monte Carlo simulation describing the spatial distribution of γ emissions in a flask. Dose rate contributed by each particle was calculated using the following equation:

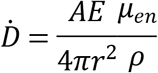

Where *A* represents the respective activity (Bq), *E* represents the respective γ energy emitted per decay (MeV) and μ_*en*_/ρ represents the mass energy-absorption coefficient (assumed to be 0.05 to represent cells). The final dose was determined in Gy through multiplying the average dose rate for all particles in the flask (determined with consideration to the previously defined geometry parameters through Monte Carlo simulation) by the time exposed to Ra-226 (9 days or 7.8 × 105 seconds).

**Fig 1.**
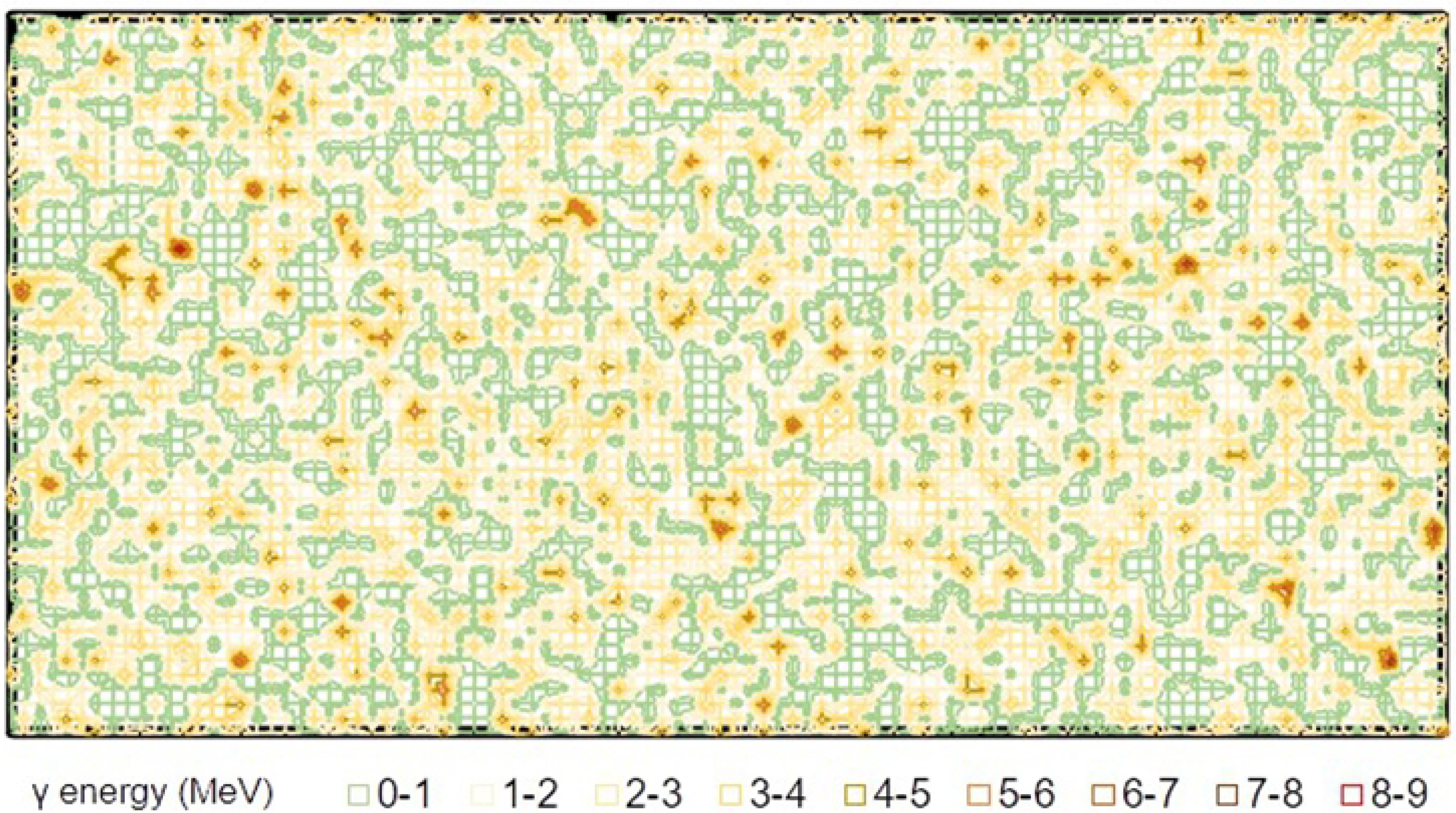
Spatial Distribution of γ-emissions. One iteration of the Monte Carlo simulation determining the spatial distribution of Ra-226 particles emitting gamma energy per emission (500 mBq/ml shown). The gamma energy is shown in MeV.

### Curve modelling

Survival fractions/plating efficiencies of cells are determined as cloning efficiency observed through staining: the fraction of colonies formed from the 500 cells plated. Residual survival fractions were calculated at each observed interval and for each dose through following the recorded cell numbers at the start and end of each interval, in accordance to previous delayed lethal effect assays by Mothersill et al. [11, 12]. Here, the product of the cells observed at the end of the current passage, with the total cell number at the end of the preceding passage, was divided by the initial number of cells seeded per passage corrected for plating efficiency. Finally, curves were fitted to the calculated residual survival values at each observed interval using the induced-repair equation taken from the model described by Lambin et al. [13]:

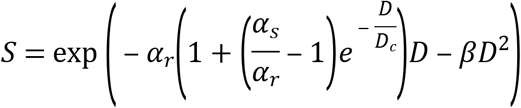

Here *a*_*r*_ describes the traditional linear-quadratic dose-response model while *a*_*s*_ describes a region of the curve showing resistance from the linear component. Curves were fit using the R Project for Statistical Computing [14] through the *nlsLM* function of MINPACK, which uses a modified Levenberg-Marquardt algorithm to perform non-linear regression. The residual sum of squares (RSS) was used to further observe the fit of the curve to empirical values, as well as verify the use of the induced-repair equation compared to the traditional linear-quadratic model.

To isolate the effect of α-particles on the survival and genomic instability of cells, this study subtracted effects observed after γ irradiation (through acute exposure to Cs-137) from mixed α and γ irradiation (through chronic exposure to Ra-226). Once the empirical data of the study is represented through curves, this is simply done through subtracting the function of one curve from the other.

## Results

### Human keratinocyte cell line (HaCaT)

Fitted curves representing the residual survival fractions for HaCaT cells show markedly different responses in directly exposed cells and their progeny with acute exposure to Cs-137 compared to those with chronic exposure to Ra-226 (Fig 2). Progenitor HaCat cells (*initial*) exposed to Cs-137 show a region of hyper-radiosensitivity (HRS) at very low doses with lower survival than would be expected by the traditional linear-quadratic model, followed by a region of increased radioresistance (IRR). The progeny of these cells continue to demonstrate such HRS/IRR behavior with further decrease in cloning efficiency thereby observing lethal mutations in those generations. In particular significant decreases in cloning efficiencies are observed in the first observation of progeny of cells (*P2, 8 population doublings*) irradiated at 0.05 Gy by 19% (*p* = 0.007), 0.1 Gy by 23% (*p* = 0.00007), 0.25 Gy by 14% (*p* = 0.02), 0.5 Gy by 15% (*p* = 0.01) and 0.75 Gy by 17% (*p* = 0.0004). Further significant decreases are observed in the second observation of progeny (*P3, 16 population doublings*) at 0.05 Gy by 17% (*p* = 0.01) and at 0.1 Gy by 14% (*p* = 0.001). The residual sum of squares (RSS) values for the initial, P2 and P3 curves are 0.0003, 0.004 and 0.002 respectively, demonstrating noticeably better fit with the induced-repair equation compared to the traditional linear-quadratic model (RSS values of 0.2, 0.09 and 0.2 respectively).

**Fig 2.**
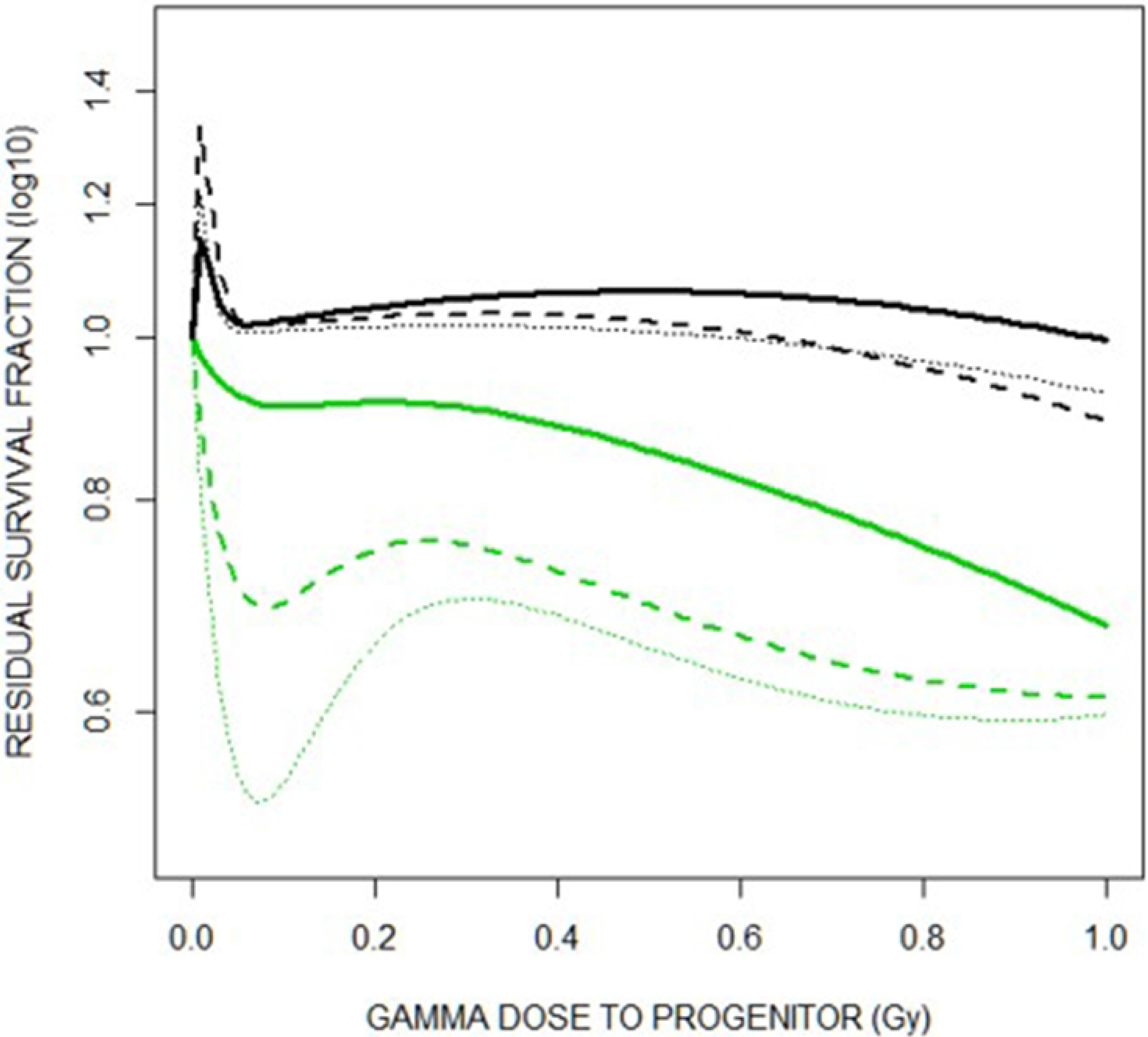
Residual Survival of HaCaT. Residual survival fractions as represented through fitted curves following the induced-repair model. Green curves represent cells exposed to Cs-137 and their progeny while black curves represent cells exposed to Ra-226 and their progeny. The darkest/solid curves represent the initial survival of the progenitors directly receiving radiation, medium/dashed curves represent the first observation of progeny (not directly irradiated), and the lightest/dotted curves represent the second observation of progeny (not directly irradiated). There were roughly 7 days between observations (time taken to reach 80-90% confluency).

In comparison, progenitor cells exposed to Ra-226 show significantly greater survival with many observations of higher cloning efficiency compared to sham irradiated control flasks (denoted as survival values greater than 100%). As such no HRS region is observed, with little to no change in survival compared to control in cells exposed to concentrations greater than 0.1 mBq/ml of Ra-226. At P2, survival of progeny in concentrations up to 10 mBq/ml of Ra-226 observe significantly higher survival compared to control, while observations at P3 show similar survival values to what was observed in the progenitors. Despite lacking an HRS/IRR region, the induced-repair equation still shows greater fit with RSS values of 0.006, 0.03 and 0.005 respectively for the initial, P2 and P3 curves (compared to 0.02, 0.08 and 0.03 respectively for the traditional linear-quadratic model), as it better matches the observed hyper increased radioresistance (HIRR) observed at very low doses.

Through subtracting the functions of fitted curves for cells exposed to Cs-137 from those exposed to Ra-226 at each interval, the relative effect of alpha exposure to the residual survival of HaCaT cells was isolated (see Table 1). Figure 3 describes the functions of the isolated effect of alpha exposure to residual survival graphically at each observation.

**Table 1.**
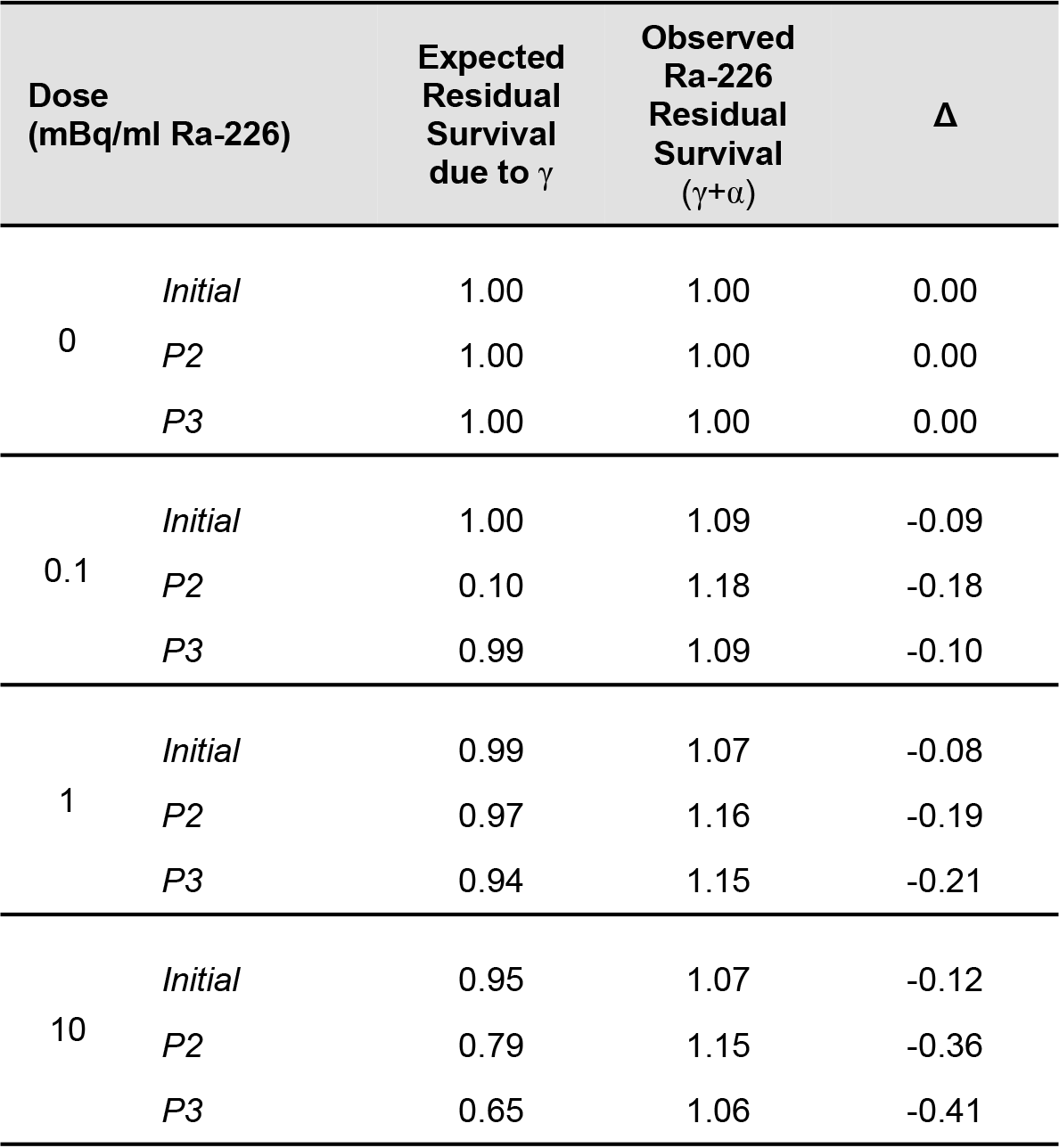
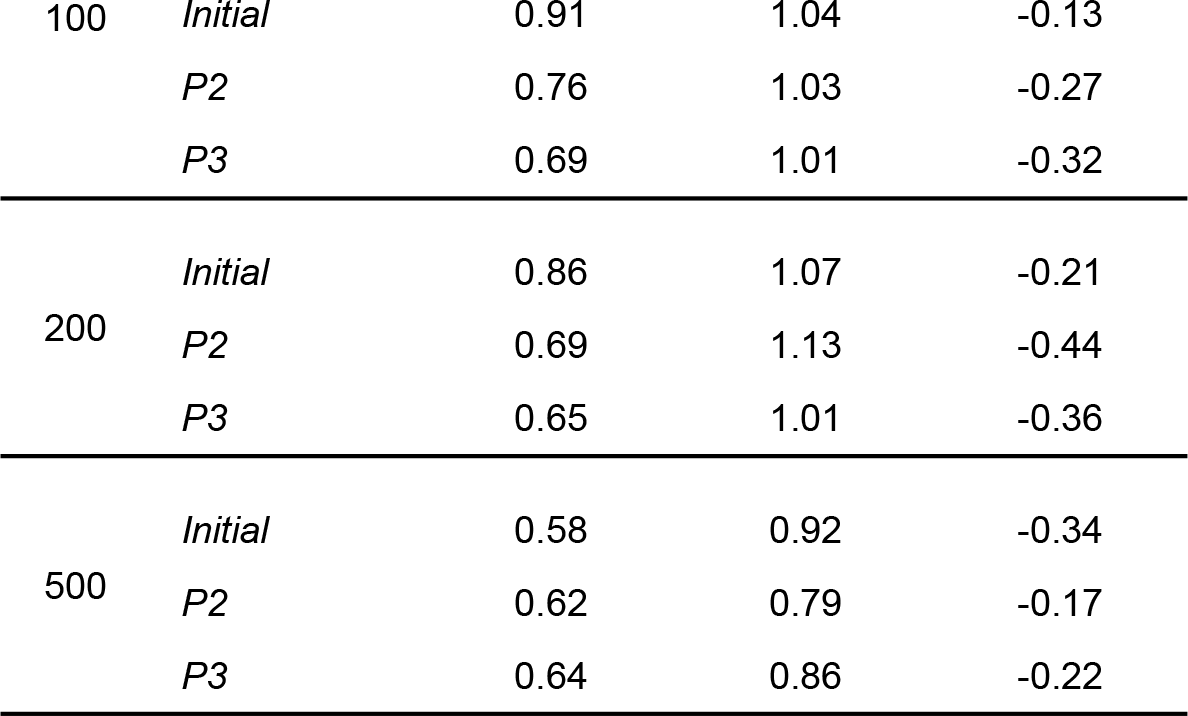
Isolating the relative effect of alpha exposure to the residual survival of HaCaT cells. At each dose, “Expected Residual Survival due to γ” was calculated using the gamma component of the dose, and the function representing residual survival of cells exposed to Cs-137. This was then compared to the residual survival observed when cells were exposed to Ra-226. The difference (Δ) is the isolated effect of alpha exposure. Note negative difference values indicate higher survival of cells in the presence of alpha particles, versus gamma exposure.

**Fig 3.**
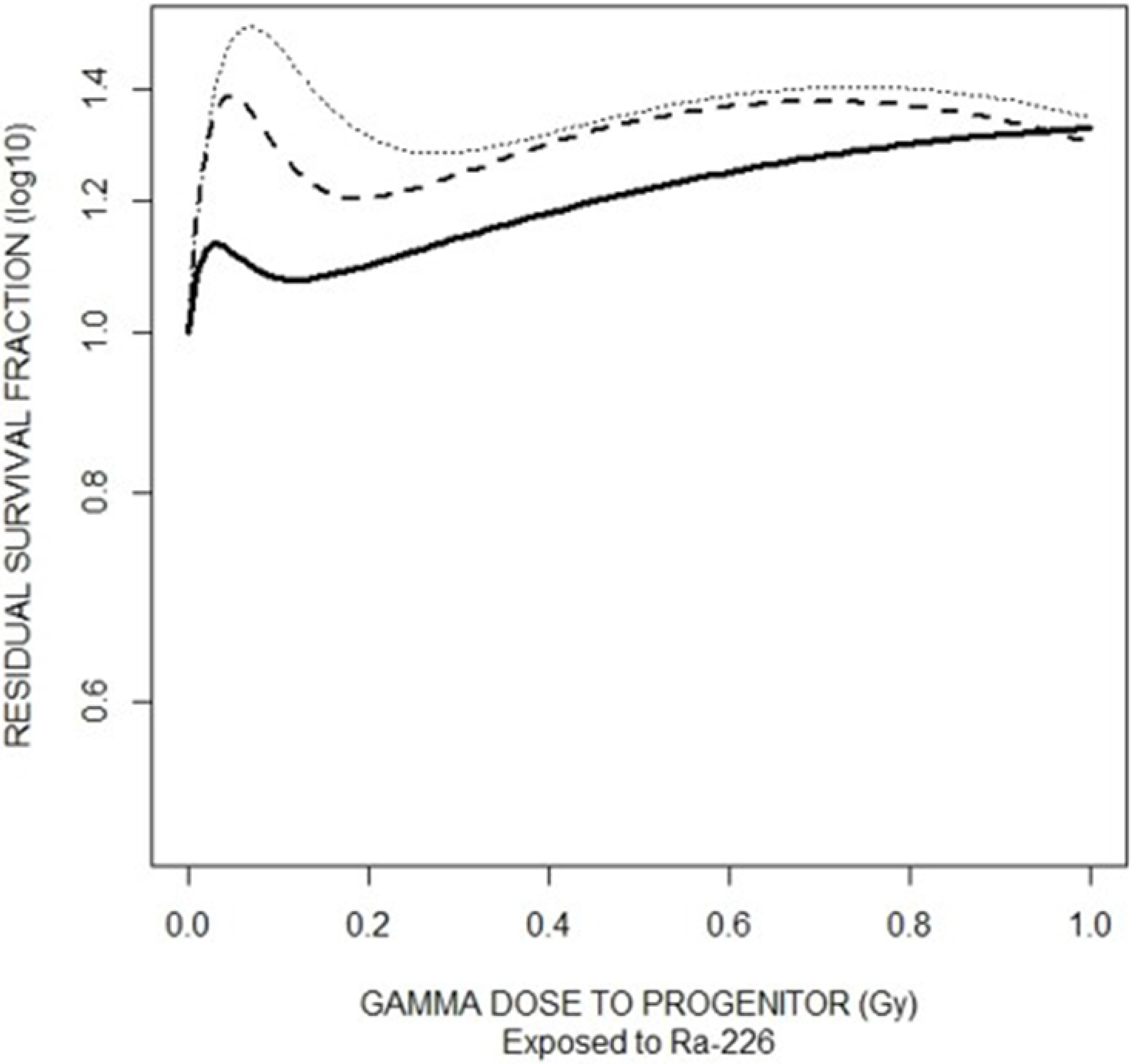
Residual Survival of HaCaT due to Alpha exposure. Calculated relative effect of alpha particle exposure on residual survival of HaCaT cells, as a function of effective gamma dose to progenitor cells. The darkest/solid curve represents the initial survival of progenitors directly receiving radiation, the medium/dashed curve represents the first observation of progeny (not directly irradiated), and the lightest/dotted curve represents the second observation of progeny (not directly irradiated). There were roughly 7 days between observations (time taken to reach 80-90% confluency).

### Embryonic Chinook salmon cell line (CHSE-214)

In contrast to the studied human cell line, there was no significant cell death observed in the directly exposed cells of the CHSE-214 fish cell line and their progeny to acute Cs-137 exposure (Fig 4). This shows an existing radioresistance when compared to human cell culture [15]. When CHSE-214 cells were exposed to Ra-226 however, progenitor cells show a marked response with decreasing cell survival following an almost linear trend with respect to dose. Residual survival observed at P2 (8 doubling periods) show increased lethal mutation however residual survival observed in the subsequent progeny at P3 (16 doubling periods) demonstrate a return of radioresistance with survival values similar to initial values.

**Fig 4.**
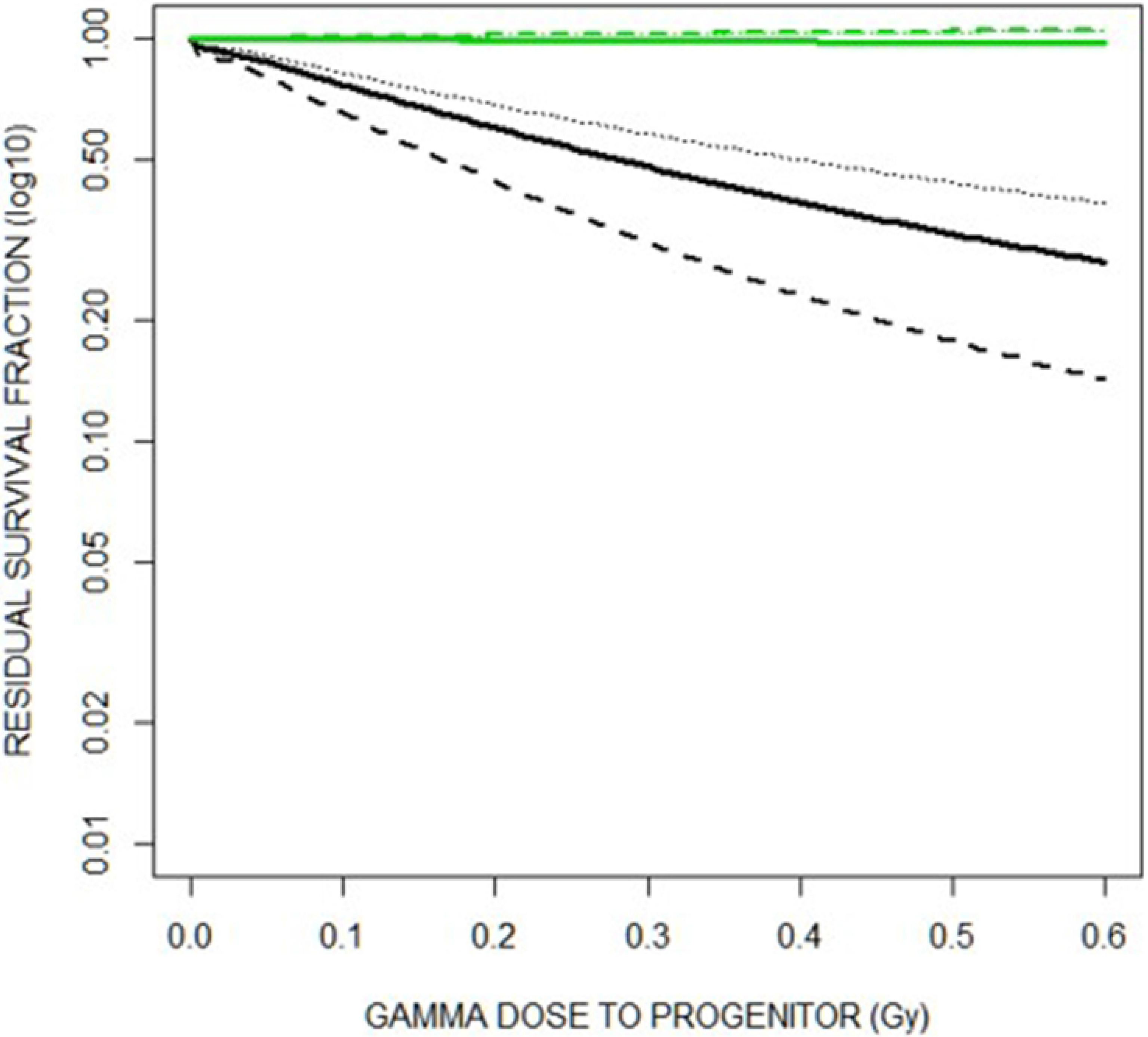
Residual Survival of CHSE-214. Residual survival fractions as represented through fitted curves following the induced-repair model. Green curves represent cells exposed to Cs-137 and their progeny while black curves represent cells exposed to Ra-226 and their progeny. The darkest/solid curve represents the initial survival of progenitors directly receiving radiation, the medium/dashed curve represents the first observation of progeny (not directly irradiated), and the lightest/dotted curve represents the second observation of progeny (not directly irradiated). Note significant overlap in residual survival of cells exposed to Cs-137 and their progeny due to minimal observed cell killing. There were roughly 40 days between observations (time taken to reach 80-90% confluency).

Using the same methodology as was done for the human cell line, the relative effect of alpha exposure to the residual survival of CHSE-214 cells was isolated (see Table 2). As exposure to gamma irradiation caused little to no effect in residual survival, the isolated relative effect of alpha exposure is significant, especially at the higher end of the low dose range. The dose dependent function for the isolated effect at each observation is shown graphically in Figure 5.

**Table 2.** Isolating the relative effect of alpha exposure to the residual survival of CHSE-214 cells. At each dose, “Expected Residual Survival due to γ” was calculated using the gamma component of the dose, and the function representing residual survival of cells exposed to Cs-137. This was then compared to the residual survival observed when cells were exposed to Ra-226. The difference (Δ) is the isolated effect of alpha exposure.

| Dose<br>(mBq/ml Ra-226) | | Expected<br>Residual<br>Survival<br>due to $\gamma$ | Observed<br>Ra-226<br>Residual<br>Survival<br>($\gamma+\alpha$) | $\Delta$ |
| --- | --- | --- | --- | --- |
| 0 | *Initial* | 1.00 | 1.00 | 0.00 |
|  | *P2* | 1.00 | 1.00 | 0.00 |
|  | *P3* | 1.00 | 1.00 | 0.00 |
| 0.1 | *Initial* | 1.00 | 0.69 | 0.31 |
|  | *P2* | 1.00 | 0.50 | 0.50 |
|  | *P3* | 1.00 | 0.66 | 0.34 |
| 1 | *Initial* | 1.00 | 0.71 | 0.29 |
|  | *P2* | 1.00 | 0.56 | 0.44 |
|  | *P3* | 1.00 | 0.73 | 0.27 |
| 10 | *Initial* | 1.00 | 0.75 | 0.25 |
|  | *P2* | 1.01 | 0.61 | 0.40 |
|  | *P3* | 1.03 | 0.81 | 0.22 |
| 100 | *Initial* | 1.02 | 0.55 | 0.47 |
|  | *P2* | 1.10 | 0.39 | 0.71 |
|  | *P3* | 1.23 | 0.64 | 0.59 |
| 200 | *Initial* | 1.02 | 0.31 | 0.71 |
|  | *P2* | 1.17 | 0.15 | 1.02 |
|  | *P3* | 1.39 | 0.45 | 0.94 |

**Fig 5.**
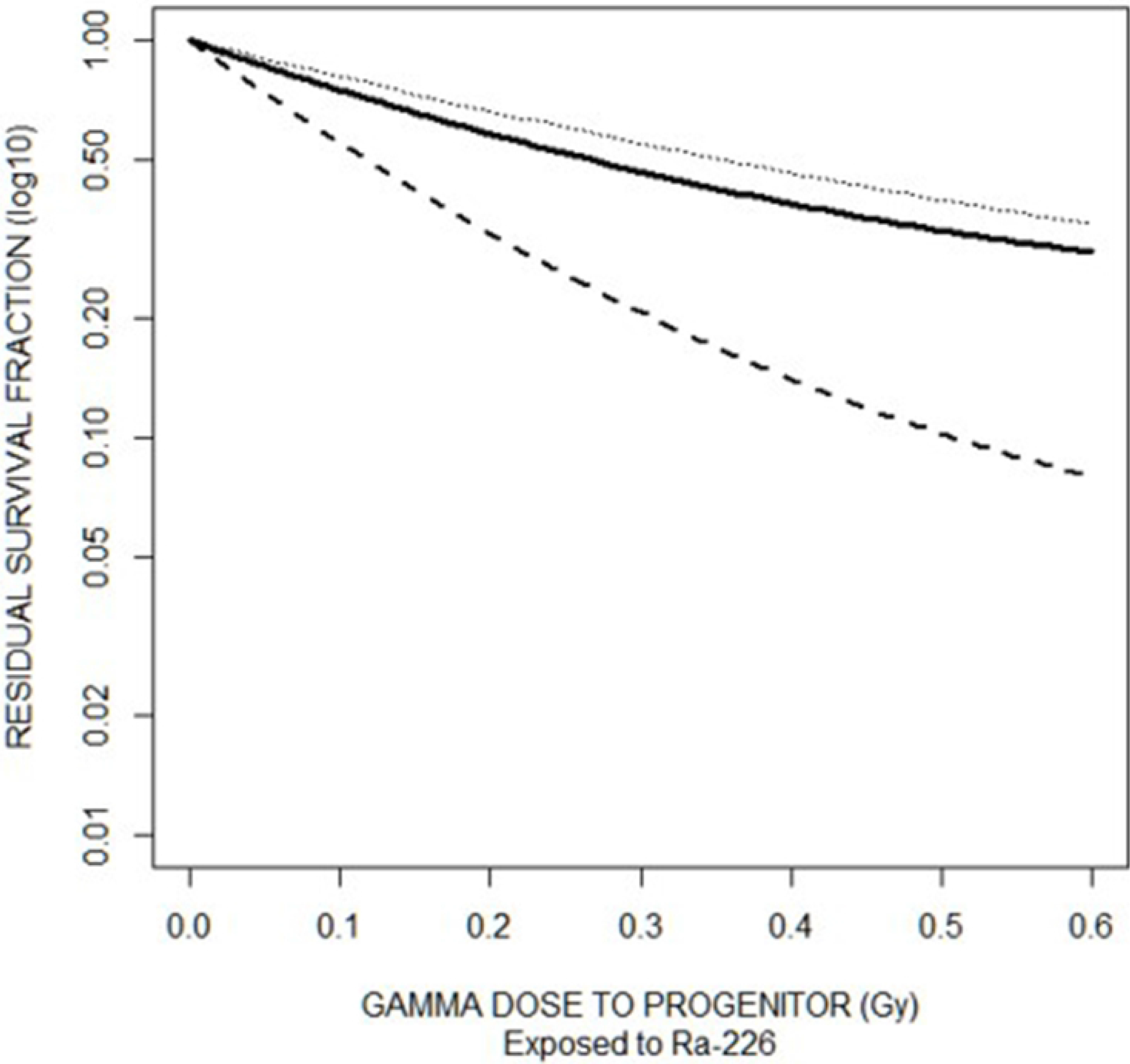
Residual Survival of CHSE-214 due to Alpha exposure. Calculated relative effect of alpha particle exposure on residual survival of CHSE-214 cells, as a function of effective gamma dose to progenitor cells. The darkest/solid curve represents the initial survival of progenitors directly receiving radiation, the medium/dashed curve represents the first observation of progeny (not directly irradiated), and the lightest/dotted curve represents the second observation of progeny (not directly irradiated).

## Discussion

At sub-lethal doses of gamma irradiation through acute exposure to Cs-137, the HaCaT cell line displayed a region of low-dose hyper-radiosensitivity (HRS) followed by increased radioresistance (IRR). In addition, lethality was observed in subsequent generations (lethal mutation phenotype) with significant decreases in cloning efficiencies observed in unirradiated progeny cells. In contrast, only radioresistance was observed in the progenitor cells exposed to Ra-226 with significantly higher survival and no observable region of HRS. Further, the observed progeny of these cells showed increased survival and lowered lethal mutation. In comparison, the results of experiments using the non-mammalian embryonic fish cell line showed the reverse of what was observed in human cell culture. Survival data following exposure to gamma irradiation confirmed existing radioresistance in the CHSE-214 cell line compared to human cell culture, with no significant lethality. However, survival data for cells exposed to Ra-226 suggested that alpha particles promoted lethality at doses otherwise known to have no significant effect.

Considering the potential for sub-lethal doses from chronic exposure to radium and its daughters found in waste products, to remnants of historic commercial and medical usage of radium (ranging from self-luminous paints to cancer treatment), the unconventional behaviors observed in both cell lines of this study have potential importance in radiological protection. Further, the presence of radium in waste reaching the ecosystem from mining and nuclear applications is important given the currently growing interest for non-human radiological protection.

The radioprotective quality of sub-lethal doses of alpha radiation that was observed in the HaCaT cell line, where cells displayed significantly lower lethality in the presence of alpha particles greatly contrasts with RBE values found in the literature. Previous *in vitro* studies of alpha radiation effects at higher doses compared to this study have all consistently demonstrated a significantly higher biological effect of alpha particles relative to photons, with values ranging from <2 for the induction of double strand breaks, to 3.5-10 for cell lethality and transformation in different cell lines, to >25 for other endpoints assessed [16]. Research observing HRS/IRR behaviors suggest the activation of cell cycle checkpoints for increased cell repair, etc. as a possible mechanism for radioresistance [17]. Considering only radioresistance was observed in the presence of alpha radiation, the results suggest an ultra-low dose of alpha particles produces a sufficient level of genomic instability to activate the previously mentioned cell cycle checkpoints, inducing radioresistance. This effect was non-linear with dose with marked reduction as dose increased to the progenitor. The results observed in the CHSE-214 cell line on the other hand is in line with currently accepted descriptions on the effect of alpha particles at high doses. Here, the concentration of damage events is said to exceed a threshold at which effective repair becomes difficult [18]. Further lethality is seen in progeny as a de novo appearance of non-clonal lethal mutations, indicative of genomic instability. However, this decreases with subsequent generations suggesting the ability for existing damage repair mechanisms eventually to counteract the heritable susceptibility to lethal damage.

The results of the study support the need to consider dose-dependence when describing the relative biological effect of different radiation qualities. Overestimation of the biological effect of sub-lethal exposure to radium in humans can result in unnecessary psychological stress and limit productivity in industry. In addition, the results of the fish cell line experiments confirm the need to be aware of species differences, confirming that protection for humans would not inherently protect ecosystems and non-human biota.

It should be noted however that the observed *in vitro* results cannot simply be translated to *in vivo* effects without further research. For example, while there is evidence for heritable NTE through *in vitro* and non-human studies, there has been no evidence for radiation-induced hereditary effects observed in epidemiological studies of human populations exposed to ionizing radiation [19]. In addition, further research needs to be done to isolate the effect of dose rate on sub-lethal exposure to high-LET radiation, as differences in time for cell repair can affect the level of radioresistance observed.

## Conclusion

At sub-lethal doses, survival greatly depends on repair mechanisms. Since the HaCaT cell line demonstrates hyper-radiosensitivity to gamma energy at low doses, high-LET alpha particle radiation may be able to produce sufficient genomic instability to induce radioresistance. In such instances, the ratio of relative biological damage caused by alpha exposure is significantly lower than an equivalent dose of gamma energy alone, and as such a lower radiation weighting factor might be considered. However, while the CHSE-214 cell line demonstrates increased radioresistance to gamma energy, the concentrated nature of energy deposited causes increased lethality when exposed to alpha particles. These cases would suggest a higher radiation weighting factor, similar to what is currently recommended. Further study is required to isolate the effect of dose-rate at sub-lethal doses. In addition, further consideration is required to translate the observed *in vitro* results to *in vivo* effects. As alpha-emitters are commonly found in industrial applications, the environment, as well as released in nuclear incidents, this knowledge would be particularly meaningful for risk management and radiation protection of human and non-human biota to low-dose high LET radiation.

## Acknowledgments

The work was funded by the Natural Sciences and Engineering Research Council (NSERC) of Canada in the form of a Collaborative Research and Development Grant (Grant No. CRDPJ 484381-15).

